# Sex-dependent noradrenergic modulation of premotor cortex during decision making

**DOI:** 10.1101/2022.12.06.519304

**Authors:** Ellen M. Rodberg, Carolina R. den Hartog, Emma S. Dauster, Elena M. Vazey

**Affiliations:** Neuroscience and Behavior Program and Department of Biology, University of Massachusetts Amherst MA

## Abstract

Rodent premotor cortex (M2) integrates information from sensory and cognitive networks for action selection and planning during goal-directed decision making. M2 function is regulated by cortical inputs and ascending neuromodulators, including norepinephrine (NE) released from the locus coeruleus (LC). LC-NE has been shown to modulate the signal to noise ratio of neural representations in target regions prior to decision execution, to increase the salience of relevant stimuli. Using rats performing a two-alternative forced choice task after administration of a β adrenergic antagonist (propranolol), we show that action planning in M2 is mediated by β adrenergic signaling. Loss of β adrenergic signaling results in failure to suppress irrelevant action plans in M2 that disrupts decoding of cue related information, delays decision times, and increases trial omissions, particularly in females. Furthermore, we identify a potential mechanism for the sex bias in behavioral and neural changes after propranolol administration via differential expression of β2 receptors across sexes, particularly on local inhibitory neurons. Overall, we show a critical role for β adrenergic signaling in M2 during decision making by suppressing irrelevant information to enable efficient action planning and decision execution.

---

Successfully navigating through an environment requires internally guided behaviors and the use of external cues to make decisions and drive goal-directed actions. Decision-making is a complex and multifaceted process, many aspects of which can be interrogated in rodents. Critical components of the decision process including detecting and encoding stimuli, forming and maintaining action plans, preparing and executing an action, and comparing predicted outcomes to the actual results of that decision. One key brain region with strong decision related activity is secondary motor cortex (M2) in rodents, analogous to Brodmann’s area 6, supplementary motor area (SMA), and pre-SMA in primates. M2 is interconnected with both sensory, motor, and cognitive brain regions (Hoover & Vertes, 2007; Jeong et al., 2016; Reep et al., 1990; Reep & Corwin, 1999) and integrates information from these networks to facilitate planning, initiation, and execution of goal-direction actions (Barthas & Kwan, 2017; Gremel & Costa, 2013; Inagaki et al., 2018; Sul et al., 2011; Wei et al., 2019). Neurons in M2 display heterogenous activity during decision-making and individual neurons differentially encode separate cues with some driving excitation and others active inhibition (Chandrasekaran et al., 2017; Inagaki et al., 2018; Wei et al., 2019). M2 neurons display some of the earliest choice related activity in the fronto-striatal network(Sul et al., 2011), encoding goal-directed action plans with neural signatures such as cue-evoked firing, activity ramping, and sustained activation during delays (Inagaki et al., 2018; Murakami et al., 2014; Svoboda & Li, 2018).

Investigating what influences choice related activity in M2 is critical to understand the mechanics of goal-directed actions. In addition to cognitive and motor network connections, M2 is innervated by neuromodulators including norepinephrine (NE) from the locus coeruleus (LC) in the brainstem (Agster et al., 2013; Hoover & Vertes, 2007). The LC is a small cluster of NE containing neurons that project throughout the brain and are the near exclusive source of NE to the cortex, including M2 (Aston-Jones, 2004; Moore & Bloom, 1979). Causal studies have shown a direct role for phasic NE in increasing gain for task-relevant stimuli (Clayton et al., 2004; Dayan & Yu, 2006; Servan-Schreiber et al., 1990; Vazey et al., 2018; Waterhouse et al., 1998). In this way, LC-NE facilitates decision-making by increasing the salience of task-related cues and maintaining attention. Although the anatomical connectivity between LC and M2 is well documented, whether and how NE facilitates action planning for decision execution via M2 is unclear.

Importantly, there is limited evidence on whether the function of noradrenergic modulation on action planning in M2 is similar across females and males. There is strong potential for interaction in these domains as there are known sex differences in both neuromodulatory clusters that provide input to M2, including LC (Bangasser et al., 2011; Pinos et al., 2001; Valentino et al., 2012) and behavioral strategies for decision making (Chen, Ebitz, et al., 2021; Chen, Knep, et al., 2021; Orsini & Setlow, 2017; Shansky, 2018).

Here we investigated how noradrenergic tone regulates M2 function and action planning in female and male rats. We recorded neural activity from M2 and modulated noradrenergic signaling through β adrenergic receptors during a simple decision-making task, two-alternative forced-choice (2AFC). We found noradrenergic blockade decreased behavioral performance and dampened the neural representations of action plans in M2, specifically by decreasing the active inhibition of irrelevant action plans. Our results show the impact of noradrenergic blockade on performance and neural activity was greater in females than males. To identify a potential mechanism for these differences in β noradrenergic sensitivity, we characterized β1 and β2 receptor levels in M2 of male and female rats. We found increased β2-adrenergic receptor expression in females that may underlie the sex specific sensitivity to noradrenergic regulation of action planning.

## RESULTS

### β-adrenergic inhibition decreases goal-directed engagement and trial completion

We sought to examine the role of norepinephrine, specifically β noradrenergic signaling, during decision-making using a 2AFC task. Rats (female n=10, male n=9) were able to quickly learn the task and perform at stable levels (>100 trials, >70% accuracy for 3 days in a row, ~2weeks). Once stable, animals were tested with the nonselective β adrenergic antagonist (propranolol, 10mg/kg, IP) or saline while recording from M2 single units. We found that propranolol decreased engagement in a 2AFC task in a sex-specific manner, impacting females more than males. We identified a reduction in the number of trials initiated after propranolol (F _(1,17)_ = 52.31; p < 0.0001; 2way ANOVA; Figure 1A) and an effect of sex and treatment interaction (F _(1,17)_ = 5.194; p = 0.0359; 2way ANOVA). Propranolol decreased initiation in females by 63% from 144.00 ± 11.46 to 52.90 ± 7.14 (p<0.0001) trials. Propranolol decreased trial initiation to a lesser extent in males (37%) from an average of 128.33 ± 9.14 to 80.89 ± 11.85 trials after propranolol (p=0.0066). Premature well exit prior to cue presentation was not affected by propranolol in either sex (means range from 13%-26%; F_(1,17)_ = 0.02224; p=0.8832; 2way ANOVA; Figure 1B). On trials where there was a cue presentation, accuracy was decreased after propranolol administration (F_(1,17)_ = 24.12; p=0.0001; 2way ANOVA; Figure 1C) particularly in females (p<0.0001) with an effect of sex and treatment interaction (F_(1,17)_ = 8.413; p=0.0100); 2way ANOVA). The change in accuracy after propranolol was driven by an increase in the percent of omitted trials in females (treatment x sex interaction F_(1,17)_ = 14.92; p = 0.0012; 2way ANOVA). Propranolol increased omissions in females four-fold from 7.73% ± 2.07% to 29.06% ± 4.59% (p < 0.0001; Figure 1D). In males, omission rate was not significantly increased after propranolol 4.13% ± 0.87% vs. 6.57% ± 1.31% (p=0.7517). It is possible that some changes in behavioral performance were a result of the non-selective or peripheral effects of β adrenergic antagonism that reduce activity across the board. We examined potential non-specific motor changes by looking at non goal-directed lever presses during the intertrial interval (ITI). Although propranolol decreased task participation in active trials, it resulted in increased intertrial interval (ITI) lever presses in both sexes by approximately 150% (F _(1,14)_ = 21.33; p = 0.0004; REML ANOVA; Figure 1E). This increase in ITI presses after propranolol indicated that goal-directed actions, rather than general lever press action was primarily impacted by β adrenergic blockade.

**Figure 1.**
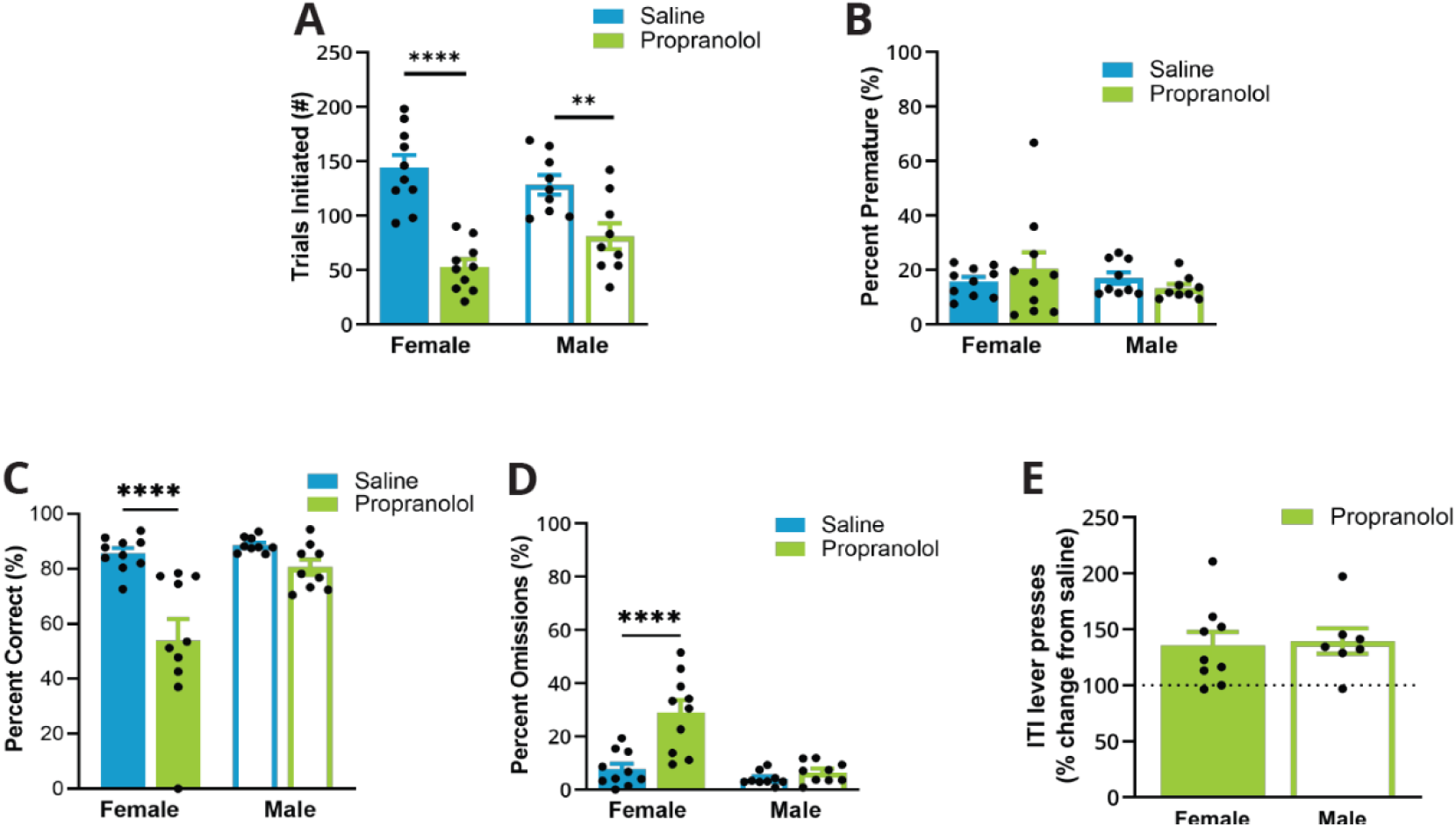
Propranolol decreased goal-directed task participation, particularly in females. (A) Propranolol (green) decreased the number of trials initiated during a 2AFC task in females (p<0.0001) and males (p=0.0066) compared to saline (blue). (B) Propranolol did not alter the percent of trials that animals left the well prior to cue presentation (premature) in females or males. (C) Propranolol decreased overall accuracy in females (p<0.001) but not males (p=0.3336). (D) The percent of trials that a response was omitted was significantly increased only in females after propranolol (p<0.0001). (E) Altered task performance was not a result of decreased motor output or lever pressing ability, measured by the ratio of lever presses made during the intertrial interval (ITI) to the total lever presses (lever press index). Compared to saline, males and females increased their lever press index by 150% after propranolol. * Indicates a significant effect compared to saline * (p<0.05); ** (p<0.01); *** (p<0.001); **** (p<0.0001).

### Propranolol does not impact the basal properties of M2 neurons

To understand the neural basis of propranolol driven changes in goal-directed decision making we recorded from M2 during this task. We collected 347 well isolated single units (female n=159, male n=188) from deep layers of M2 across 14 rats (female n=8, male n=6) after saline vehicle or propranolol (Figure 2A, B). Within a session, M2 displayed heterogeneous event related activity. We identified two primary task-related activity drivers: cue onset (‘Cue’) and lever press response (‘Lever Press’) (Figure 2C). Action planning and choice signals in M2 were well represented in both sexes, however cue and/or lever press related activity were less common in female subjects. In females, the percent of M2 units with task locked modulation was 77.55% after saline and 72.13% after propranolol. In males, 93.46% of units demonstrated task locked modulation after saline and 85.19% after propranolol. This base sex difference in task evoked M2 activity was not impacted by β-adrenergic signaling (female χ^2^(1, n=159) =0.5966, p=0.4399; male χ^2^((1, n=188)=1.236, p=0.2662; Chi-squared test). Only units with cue or lever press task related activity were used for further analysis (saline n=176, female n=76, male n=100; propranolol n=116, female n=44, male n=72).

**Figure 2.**
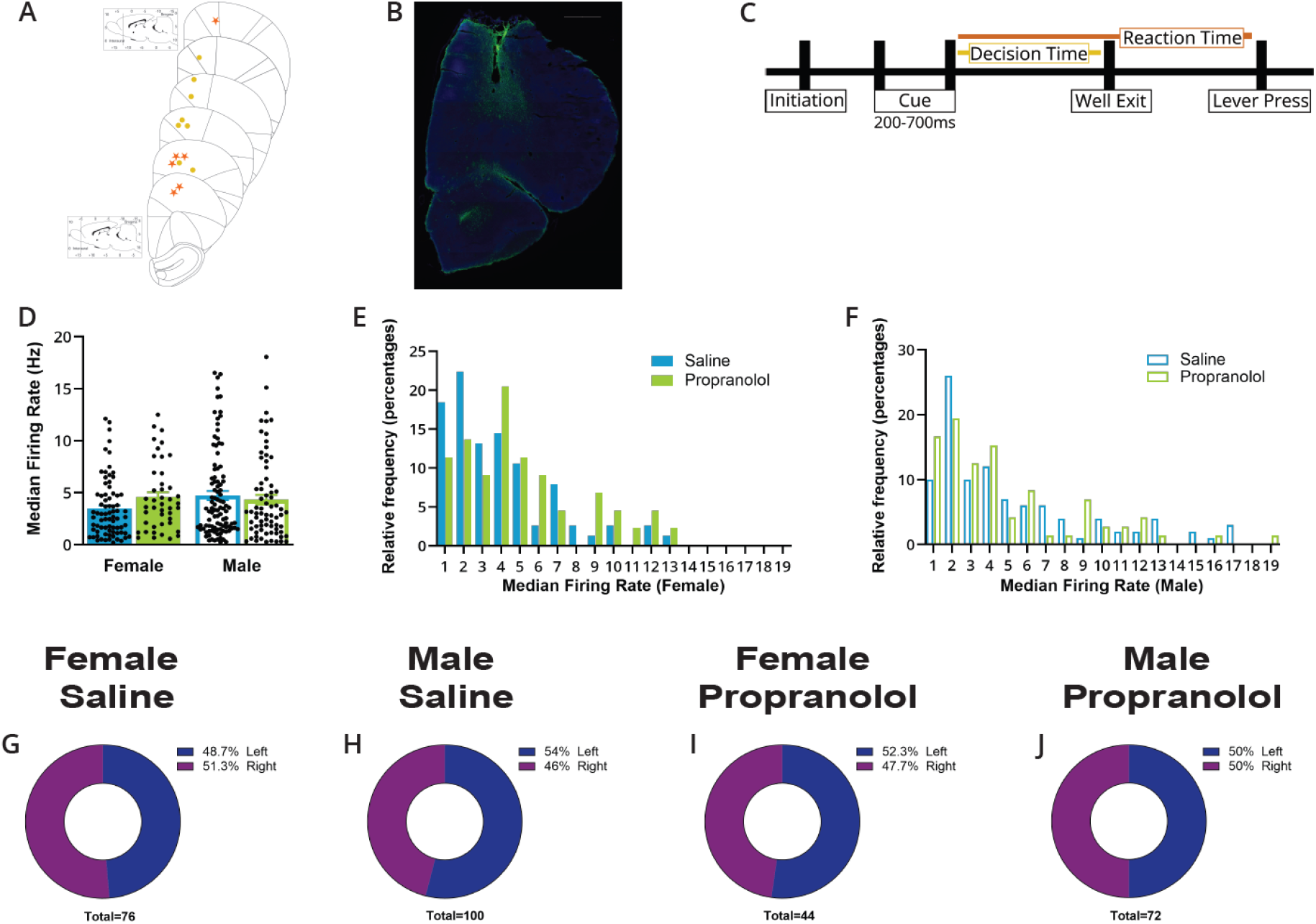
Basal properties of M2 units are not altered by propranolol. (A) Placements of the center of electrode array in M2 in females (yellow circle) and males (orange star). (B) Representative figure of histological placement of microelectrode array in M2. Tissue is stained with DAPI (blue) and GFAP (green) to identify glial scarring after electrolytic lesions. Scale bar indicates 1000μm. (C) Schematic of events in the 2AFC task. Rats self-initiated trials by breaking an IR beam in the reward well to receive one of two randomly selected cues that informed which lever would be rewarded when pressed. Cues were presented at a variable interval 200-700ms after initiation and until animals left the well, ‘Well Exit’. The time from cue onset to well exit was identified as the decision time (yellow). Animals had 5s after cue presentation to execute a decision via lever pressing, ‘Lever Press’. The time from cue presentation to lever press was identified as the reaction time (orange). Possible trial outcomes were correct, incorrect, omission (no lever press within 5s), or premature (well exit prior to cue presentation). (D) Propranolol did not alter basal firing properties (median firing rate) in females (p=0.2226) or males (p=0.7227). Median firing rate (Hz) of each unit included in electrophysiology are represented as individual points. Bar graphs represent the mean ± SEM of units from females and males after saline (blue) and propranolol (green). (E) The distribution of single unit median firing rate (Hz) in females was not significantly different after propranolol (p=0.0845; Kolmogorov–Smirnov test). Histograms show the distributions of median firing rates in females after saline (blue) and propranolol (green) in 1s bins where the value of the x axis is the maximum value in each bin. (F) The distribution of single unit median firing rate (Hz) in males was not significantly different after propranolol (p=0.7517; Kolmogorov–Smirnov test). Histograms show the distributions of median firing rates in males after saline (blue) and propranolol (green) in 1s bins where the value of the x axis is the maximum value in each bin. (G-J) Propranolol does not change the distribution of side preference displayed in M2 neural activity. Proportions of units that display a left (blue) or right (purple) side preference during the 2AFC task separated by sex and treatment. Side preference was identified as the largest Z-score of 100ms bins at the time of significantly different single unit neural activity on right and left trials (G: female saline, H: male saline, I: female propranolol, J: male propranolol). Total number of units per treatment and sex is identified at the bottom of the graphs and the values in figure legends identify the percent of units in each classification.

Across the session median firing rates in task encoding neurons were similar between sexes 3.47 ± 0.33Hz in females vs 4.75 ± 0.42Hz in males after saline, and 4.57Hz ± 0.49Hz vs 4.33Hz ± 0.47Hz after propranolol (effect of sex F _(1,288)_ = 1.318; p = 0.2520; 2way ANOVA). Propranolol did not impact the median firing rate (effect of treatment F _(1,288)_ = 0.5776; p = 0.4479; 2way ANOVA; Figure 2D) or distribution of firing rates recorded in either sex (female p=0.0845, male p=0.7517; Kolmogorov–Smirnov test; Figure 2E, 2F). As previously noted, individual M2 units exhibit side preference/laterality encoding with preferential increased firing for one cue and/or lever press side, and the same neuron undergoing active inhibition for the opposite cue/lever press (Li et al., 2016; Inagaki et al., 2018; Wei et al., 2019). The proportion of neurons preferring either left or right cues and action sequences was unaffected by sex or propranolol (female χ^2^(2, n=159) =0.7376, p=0.6916; male χ^2^((2, n=188)=1.503, p=0.4718; Chi-squared t-test; Figure 2G-J).

### β-Noradrenergic signaling is critical for suppression of irrelevant action plans in M2

We further investigated how propranolol influenced encoding of task cues in M2. Visual cues led to differential responses in individual neurons with preferred side cues driving excitatory activity, and non-preferred cues driving active inhibition (representative neuron, Figure 3A). The latency to cue discrimination across trials (when each unit’s activity was significantly different on left vs. right cued trials) was consistent between sexes (sex F_(1,201)_ = 1.017; p=0.3144; 2way ANOVA) and delayed after propranolol treatment (treatment F_(1,201)_ = 6.757; p=0.0100; 2way ANOVA; Figure 3B, 3C). Further, propranolol significantly increased the proportion of units that failed to discriminate left vs right cues in females from 26.32% to 47.73% (p=0.0273; Fisher’s exact test) and males from 15.00% to 43.06% (p<0.0001; Fisher’s exact test), severely disrupting overall neural signatures of action plans (Figure 3B, 3C, leftmost bars).

**Figure 3.**
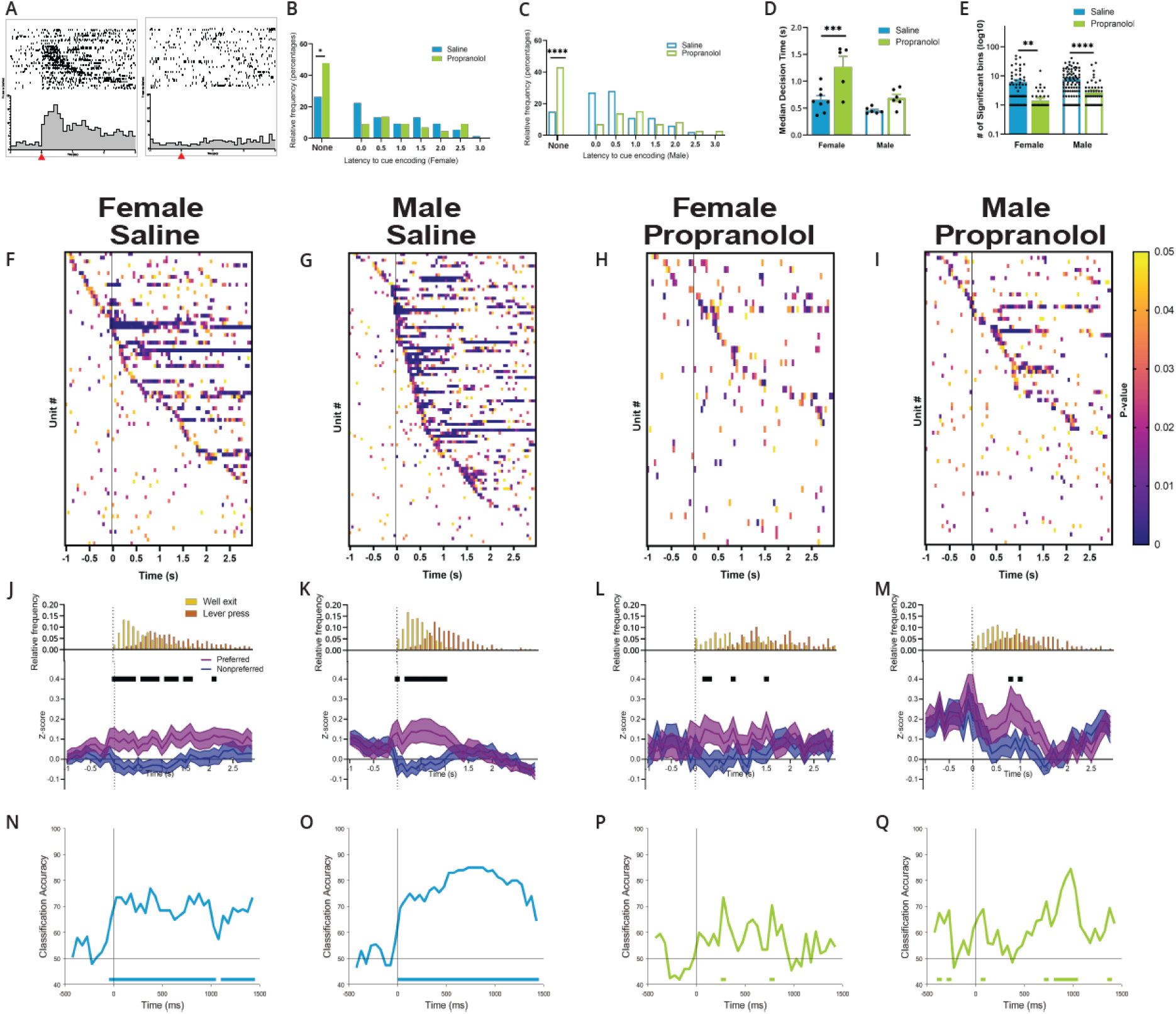
Propranolol decreases active inhibition of nonpreferred action plans. (A) Differential responses to opposing cue presentations. Raster (above) and perievent histogram of a representative neuron aligned to cue onset (t=0s, 100ms bins) from 1s prior cue onset until 3s after cue onset. Leftmost graph shows a single units’ response to left cue presentation and the rightmost graph shows that same units’ response to right cue presentation. This representative neuron was classified as left preferring, exemplified by a sharp and sustained increased in neural activity at left cue presentation but not right cue presentation. Red arrow on the x axis identified cue onset at time=0s. (B) Propranolol increases the proportion of units that do not discriminate left vs right cues in females. Latency to show differential activity on left vs right cued trials was unchanged (500ms bins; p=0.8126; Kolmogorov-Smirnov test). Leftmost bars indicate units that never encode cue directionality (p=0. 0.0273; Fisher’s exact test) which was increased by propranolol. (C) Propranolol increases the proportion of units that do not discriminate left vs right cues in males. Latency to show differential activity on left vs right cued trials also changed after propranolol (500ms bins; p=0.0045; Kolmogorov-Smirnov test). Leftmost bars indicate units that never encode cue directionality p=0. <0.0001; Fisher’s exact test) which was increased by propranolol. (D) The latency to behaviorally discriminate cues and select an action, measured by the time from cue onset to well exit (decision time), differed by sex and propranolol. Propranolol significantly increased the median decision times in females (p=0.006) but not males (p=0.2172). (E) Propranolol decreased the duration of task relevant neural activity around cue onset (100ms bins; −1s to 3s) in both females (p=0.0021) and males (p<0.0001). (F-I) Heatmaps displaying the p-values of comparisons of single unit neural activity on right vs left trials around cue onset show propranolol’s effect on the strength and duration of cue encoding and the resulting action planning. Each heatmap is separated by sex and treatment (F: saline female, G: saline male, H: propranolol female, I: propranolol male) aligned to cue onset (100ms bins; −1s to 3s). Significant p-values are represented in colored boxes from 0.05 (yellow) to approaching 0 (purple). (J-M) Propranolol decreases the active inhibition of neural activity on nonpreferred cued trials in females and males. Neural population activity between preferred and nonpreferred trials after cue onset corresponds with behavioral measures of action selection. Population neural data split by preferred and nonpreferred cue onset separated by sex and treatment (J: saline female, K: male saline, L: propranolol female, M: propranolol male). (J,K) Both sexes show task-related activity began shortly after cue onset and remained patterned, corresponding with time to well exit (represented in histograms above Z-scored graphs, yellow) after saline. Neural activity on preferred (purple) and nonpreferred (blue) trials show sharp and sustained increases and decreases, respectively, from cue onset until well exit. (L) Propranolol decreased the difference between preferred and nonpreferred activity in females by decreasing task related activity in both populations. (M) Propranolol blocked the nonpreferred decrease in males, increasing the noise represented in task-related activity. Graphs show mean ± SEM of Z-scored activity on preferred (purple) and nonpreferred (blue) trials. Black bars above the graphs indicate significantly different population activity. Histograms above the Z-scored population activity show the distribution of timepoints for well exit (yellow) and lever press (orange) relative to cue onset. (N-Q) Decoding cue type from M2 neural activity was demolished after propranolol treatment in females and males. Graphs show the result of permutation testing of a trained decoder for each group (N: female saline, O: male saline, P: female propranolol, Q: male propranolol). Classification accuracy for male and female saline (blue) groups were significantly above chance starting at cue onset and persisting for the remaining 1.5s. Classification accuracy decreased in both female and male groups after propranolol (green) with almost no performance above chance. Graphs show classification accuracy of decoding model trained with equal number of neurons (40) and trials (10/cue type) in 150ms bins. Bars on the bottom of each graph show time bins where the classification accuracy was significantly above chance.

This reduced discriminability of cues in M2 was reflected behaviorally by an increase in median decision time (time from cue onset to well exit), after propranolol (F_(1,21)_ = 17.63; p = 0.0004; 2way ANOVA; Figure 3D). There was an effect of sex on the impact of propranolol on decision time (F_(1,21)_ =14.74; p = 0.0010; 2way ANOVA) where median decision time in females showed the greatest disruption after propranolol (p=0.0006; Sidak’s multiple comparisons test). The distribution of decision times was rightward shifted and flattened in both sexes showing delayed decision processing and increased variability after propranolol (female F_(1,1024)_ =78.30, p<0.0001; Brown-Forsythe; male F_(1,862)_ =35.57, p<0.0001; Brown-Forsythe). Propranolol almost doubled median decision time in females from 653.7ms ± 79.3ms to 1265ms ± 198.3ms and increased median decision time in males from 454.2ms ± 23.23ms to 689.9ms ± 68.89ms.

We saw a reduction of sustained cue encoding in both sexes after propranolol (F_(1,288)_ = 28.20; p < 0.0001; 2way ANOVA; Figure 3E). Propranolol reduced the average number of 100ms time bins single neurons significantly discriminated between left vs right cues in females from 6.07 ± 1.13 to 1.41 ± 0.37 (p=0.0021) and males from 7.62 ± 0.82 to 2.65 ± 0.57 (p<0.0001). The reduction in cue discrimination and action planning signals were seen in both individual units (Figures 3F–3I) and across the population (Figures 3J–3M). Across the population, post cue differential encoding was largely sustained over a 1.5s epoch in females and 1.1s in males on saline sessions (Figures 3J–3K). The duration of sustained cue encoding in M2 encompassed the timing of action planning (well exit) and execution (lever press) when comparing to behavioral response characteristics (Figures 3J–3M upper inserts). After propranolol, cue representations were only present for short 100-200ms bursts and was not widely apparent in individual units (Figures 3H–3M) or the population signal (Figures 3L–3M). The loss of cue discrimination in M2 was primarily driven by a loss of active inhibition for the nonpreferred cue. The suppression of irrelevant action plans was severely dampened after propranolol introducing significant noise into decision-making process in M2.

To verify whether relevant task-related information remained in M2 after propranolol we used a neural decoding toolbox(Meyers, 2013) to compare population decoding accuracy. We sparsely trained a linear support vector machine classifier under each condition (female saline, male saline, female propranolol, male propranolol). We then compared decoding accuracy for each group to shuffled data and identified time bins where the decoding results were greater than chance based on a permutation test. We found, given the same training parameters, decoding accuracy was decreased in female (Figures 3N, 3P) and male (Figures 3O, 3Q) populations after propranolol administration compared to saline. This result shows that without β adrenergic signaling, M2 neurons no longer adequately represent information about cue type and therefore no longer appropriately encode action plans.

Overall, these data show that cue encoding and the resulting action plans within M2 are dependent on noradrenergic activity. Loss of β noradrenergic signaling disrupts suppression of irrelevant action plans leads to a failure to engage appropriate goal-directed actions, increasing decision times (Figure 3D), increasing omitted trials (Figure 1D), and reducing task engagement (Figure 1A).

### Propranolol disrupts action plan representation in M2 leading up to decision execution

We next analyzed the neural data around action execution by aligning neural activity to lever press responses on completed trials. Again, single units showed heterogenous activity around critical task epochs and distinct side preferences for opposite lever press responses (representative neuron, Figure 4A). For completed lever presses propranolol did not alter the latency of action selection in M2 prior to lever press in females (saline −1.41s ± 0.12s, propranolol −1.64s ± 0.15s) or males (saline −1.45s ± 0.09s, propranolol −1.54s ± 0.13s) (F_(1,215)_ = 1.713; p = 0.1920; 2-way ANOVA; Figure 4B,C). Propranolol did not change the proportion of units that expressed an action plan prior to decision execution in females (saline=75%, propranolol=65.91%; p=0.3009; Fisher’s exact test) but slightly decreased the proportion of units expressing an action plan in males (saline=83%, propranolol=69.44%; p=0.0431; Fisher’s exact test). Unlike cue aligned data (Figure 3B, 3C) where subsequent actions may have been omitted, lever press completion inherently required a prepotent action plan in M2 which was still evident under propranolol despite the occurrence of these events throughout the session being greatly reduced (Figure 4B, 4C).

**Figure 4.**
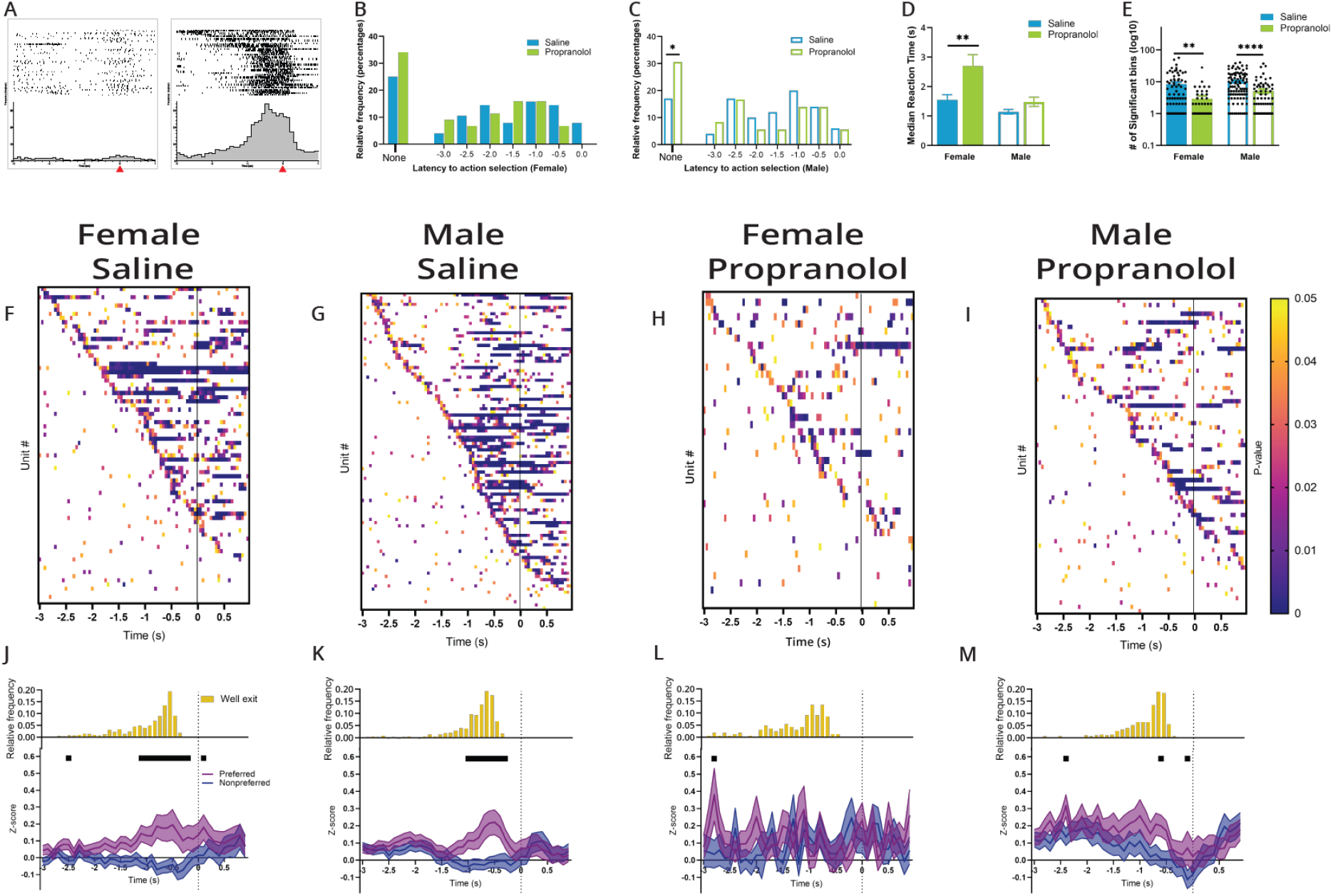
Propranolol decreases action plan representation at decision execution, increasing reaction times. (A) Differential responses to preferred and non-preferred lever press. Raster (above) and perievent histogram aligned to lever press response (at t=0s,100ms bins) from 3s before lever press to 1s after lever press. Leftmost graph shows a single units’ response to a left lever press and the rightmost graph shows that same units’ response to a right lever press. This neuron was classified as right preferring, exemplified by a sustained increased in neural activity leading up the right lever press but not left lever press. Red arrow on the x axis identified lever press at time=0s. (B) In females, propranolol (green) did not change the proportion of M2 units that encode action selection (p=0.3009) or the latency to discriminate (p=0.4394) upcoming lever press response on completed trials from saline (blue). Latency distribution (500ms bins) from 3s prior to lever press with the proportion of units that never discriminate lever press in the leftmost bars. (C) In males, propranolol (green) slightly decreased the proportion of M2 units that encode action selection (p=0.0431) but had no effect on the latency to discriminate (p=0.8379) upcoming lever press response on completed trials from saline (blue). Latency distribution (500ms bins) from 3s prior to lever press with proportion that never discriminate lever press in the leftmost bars. (D) Behaviorally, median reaction times, measured by time from cue onset to decision execution, were significantly altered by propranolol and sex. Propranolol significantly increased median reaction times in females (p=0.0015) but not males (p=0.4606). (E) Propranolol significantly decreased representation of distinct lever press related activity from 3s before to 1s after lever press (100ms bins). β noradrenergic blockade reduced the duration of action planning in both females (p=0.0023) and males (p=0.0004). (F-I) Heatmaps of single unit neural activity on right vs left response trials around lever press show propranolol’s effect on the strength and duration of action planning. Each heatmap is separated by sex and treatment (F: saline female, G: saline male, H: propranolol female, I: propranolol male) aligned to lever press (3s before to 1s after lever press) showing the 100ms bins where single unit activity on right and left response trials were significantly different. Significant p-values are represented in colored boxes from 0.05 (yellow) to approaching 0 (purple). (J-M) M2 dissociates actions from well exit through action execution (leverpress). Propranolol decreased the active inhibition of action plans for the opposing non-preferred lever in both sexes, and goal-directed action plan for the target (preferred) lever in females. Population neural data split by preferred (purple) and nonpreferred (blue) lever press trials separated by sex and treatment (J: saline female, K: male saline, L: propranolol female, M: propranolol male). (J,K) Both sexes showed task-related activity began shortly (~1s) prior to lever press and remained until action execution, corresponding with well exit times (histograms above Z-score graphs) after saline. Neural activity on preferred and nonpreferred trials show sharp and sustained increases and decreases, respectively, in activity prior to lever press execution. (L) Propranolol decreased the difference between preferred and nonpreferred activity in females by demolishing task related activity in both populations. (M) Propranolol decreased the difference between preferred and nonpreferred activity in males by decreasing inhibition on nonpreferred trials, increasing the noise represented in task-related activity. Graphs show mean ± SEM of Z-scores, black bars above the graphs show timepoints were the population activity of preferred and nonpreferred activity were significantly different. Histograms above the Z-scored population activity show the distribution of well exit times, respective to lever press (yellow).

Behaviorally, median reaction times, measured by the time from cue onset to decision execution (lever press), were significantly altered by propranolol (F_(1,21)_ = 12.82; p = 0.0018; REML ANOVA) and sex (F_(1,21)_ = 15.37; p = 0.0008; REML ANOVA; Figure 4D). Propranolol delayed median reaction times from 1.55s ± 0.17s to 2.70s ± 0.38s in females (p=0.0015) but did not alter median reaction times in males, from 1.14s ± 0.08s to 1.48s ± 0.15s (p=0.4606). Reaction time distribution within both sexes shifted rightward and broadened with increased variance after propranolol compared to saline (female p<0.0001, male p<0.0001; Kolmogorov-Smirnov test). This disruption in reaction time may be related to impairments in sustained action plan signaling in M2. Although action plans were identifiable in some neurons after propranolol, across M2 sustained representation was disrupted by β noradrenergic blockade (F_(1,288)_ = 24.27; p < 0.0001; 2way ANOVA; Figure 4E). Propranolol reduced the average number of 100ms bins single neurons discriminated between left vs right lever press responses in M2, from 8.37 ± 1.25 to 2.93 ± 0.69 bins (p=0.0023) in females and 10.17 ± 0.94 to 5.69 ± 0.823 (p=0.0004) in males. Individual unit activity leading up to lever press/decision execution exemplifies the disruption propranolol had on action selection (Figure 4F–4I). Across the population, action plans were represented in M2 around 1s before lever press through decision execution under vehicle conditions (Figure 4J, 4K). Propranolol primarily disrupted active inhibition of irrelevant action selection in M2 activity, reducing the ability to maintain action plan representation in M2, delaying lever press responses (Figure 4L, 4M). Thus, the ability to select and maintain an action plan in M2 is greatly facilitated by β adrenergic signaling. The disruption in the ability to maintain an action plan selection in M2 was most severe in females who also showed behaviorally the greatest increases in reaction times.

### Females show greater β2 adrenergic receptor density in M2

We profiled adrenergic receptor populations within M2 using RNAscope (Figure 5A) to determine whether biased adrenergic signaling may contribute to the sex-based differences in propranolol’s modulation of behavior and neural activity. We quantified β1 and β2 adrenergic receptors as these subtypes are prominently expressed in frontal cortical regions (Nicholas et al., 1991; Rainbow et al., 1984; Wanaka et al., 1989). Further, propranolol is selective for β1 and β2 but not β3 receptors (Baker, 2005; Schena & Caplan, 2019). We found females had higher levels of β adrenergic receptors in M2 than males (effect of sex F_(1,14)_ = 7.48; p =0.0161; 2way ANOVA; Figure 5B), particularly β2 adrenoreceptors (p=0.01; Sidak MCT). To determine whether β adrenergic modulation was influencing the output of pyramidal neurons in M2 or local interneurons we colabelled for either excitatory(vGLUT1) or inhibitory(vGAT) amino acid transporter mRNA. We found that much of the overall increase in β2 adrenoreceptor density in females was located on vGAT+ neurons within M2 (effect of cell type F_(2,28)_ = 7.02; p =0.0034; 2way ANOVA; Figure 5D). β1 adrenoreceptors were near exclusively found on vGLUT+ neurons (F_(1,14)_ = 225.5; p <0.0001; 2way ANOVA; Figure 5C). This evidence provides a potential mechanism whereby decision making in females critically relies on β2 adrenergic modulation of local interneurons to suppress irrelevant information signaling in M2 for efficient action planning.

**Figure 5.**
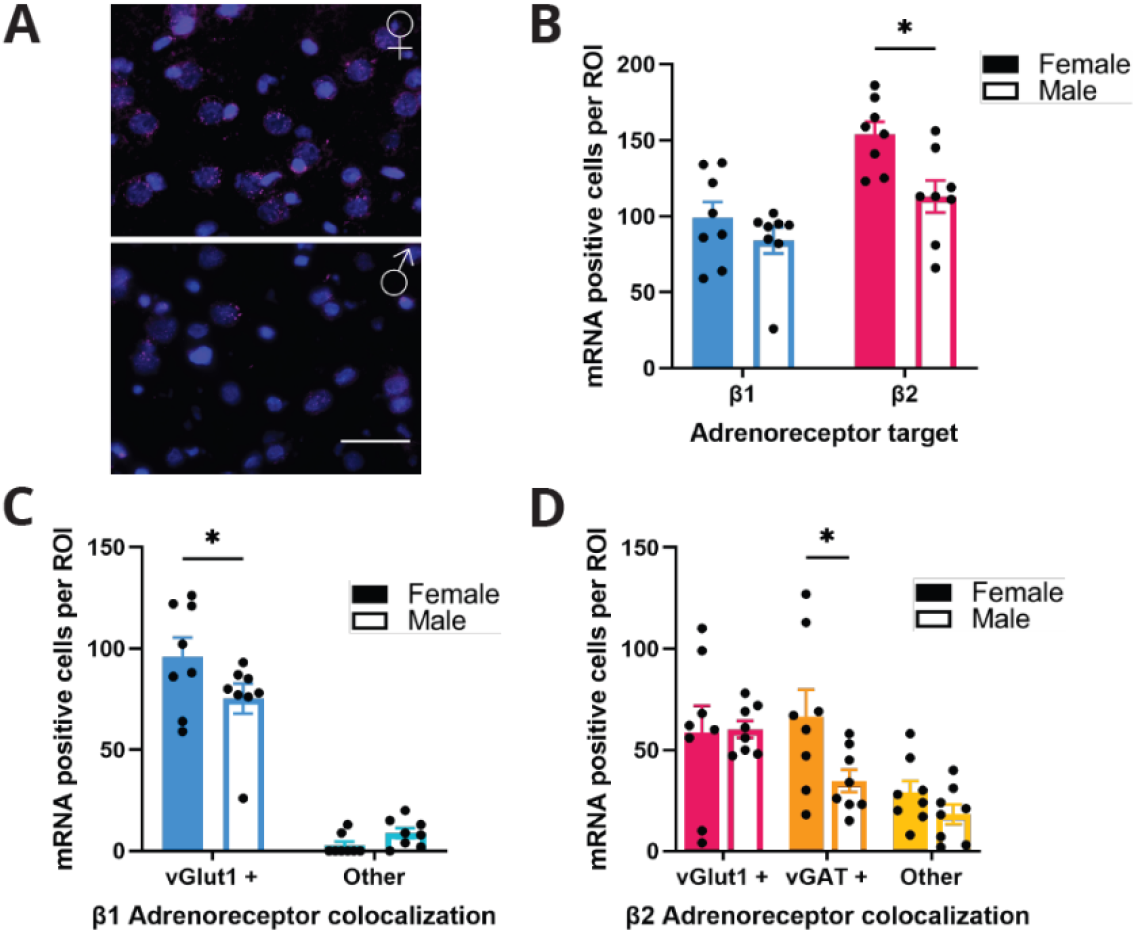
Females express more β adrenoreceptors in M2. (A) Representative figure of RNAscope in M2 of female (top) and male (bottom) rodents. β2 mRNA expression (magenta) in M2 with DAPI (blue) counterstain. Scale bar indicates 50 μm. (B) Females have a greater expression of β2 receptor mRNA (pink) in M2 compared to males (p=0.01). There was no difference in β1 receptor mRNA (blue) expression in females and males. (C) β1 receptor expression was significantly higher on vGlut1+ cells (blue) in females compared to males. There was no sex difference in the amount of β1 receptor expression on non-vGlut1+ cells (light blue). (D) β2 receptor expression was significantly higher on vGAT+ (orange) but not vGlut1+ (pink) or other (yellow) cells in females compared to males.

## DISCUSSION

How does norepinephrine impact the ability for M2 neurons to encode action plans during goal-directed decision making? Here we look at noradrenergic regulation of action planning by recording from M2 neurons under β-adrenergic blockade during a 2AFC task and characterizing β receptor expression in M2. We found that blocking β-adrenergic signaling blunted task relevant neural activity, most notably reducing suppression of irrelevant action plans. Propranolol decreased the information encoded in M2 during the 2AFC resulting in decreased task engagement and performance. The degree to which these behavioral and neural deficits occurred differed by sex where females displayed greater dependence on β-adrenergic activity for action planning. Females also showed higher levels of β-adrenergic receptor mRNA within M2, particularly on local interneurons. Results from this study suggest that noradrenergic signaling on β receptors is key to maintaining neural representation of action plans necessary for optimal performance during decision-making and provides evidence for sex differences within this mechanism.

The LC-NE system has been implicated in signal detection, altering gain of task relevant cues in sensory cortices. NE acting in these regions amplify salient cue related signals while simultaneously quieting aberrant or ‘noisy’ background activity (Clayton et al., 2004; Dayan & Yu, 2006; Servan-Schreiber et al., 1990; Vazey et al., 2018; Waterhouse et al., 1998). Our data confirms this role for NE in higher cognitive regions such as M2 and identifies that this mechanism can work specifically through β adrenoreceptors. The overall effect of blocking β-adrenergic signaling was that fewer neurons encoded task relevant information (Figure 3B&C) and for a shorter period of time (Figure 3E, 4E). This delayed formation of an action plan in M2, particularly in females, and led to increased response times (Figure 3D, 4D) and a greater rate of omissions (Figure 1D). Blocking β noradrenergic activity decreased the magnitude of (Figure 3J-M), but not latency (Figure 3B&C), of cue encoding. This maintained latency suggests that propranolol did not affect the detection of task relevant stimuli but did impact the successive information integration and resulting action plan. Using a linear classifier, we showed that cue type could be decoded from M2 neural activity after saline but decoding accuracy was lost after propranolol (Figure 3N-Q). Diminished cue encoding after propranolol necessitated a longer period of time for the formation of an action plan and thus, delayed decision execution (Figure 3D, 4D). This evidence supports a role for noradrenergic signaling in M2 – optimizing behavioral engagement and the gain of action plan representation.

Previous studies have shown M2 neurons display heterogenous activity during decision-making tasks (Chandrasekaran et al., 2017; Inagaki et al., 2018; N. Li et al., 2016; Svoboda & Li, 2018). In particular, M2 units show preferential activation for cues/actions based on egocentric location and active inhibition for opposing cues/actions (Inagaki et al., 2018; Wei et al., 2019). Our data replicated these findings and show these functions are consistent across sexes with similar basal (Figures 2D-F) and task-related (Figures 3J,K & 4J,K) activity in M2 neurons between females and males. In the current study, suppressing β adrenergic signaling did not alter baseline firing properties in either sex but caused targeted deficits in neural representations of task related information. Our findings identify a specific role for β-adrenoreceptors in selectively regulating the accurate representation of action plans in M2 during goal directed decision making.

Distinctions between the cognitive effects of β1 and β2 receptor activity in the brain have identified that β2 agonism improves working memory (Ramos et al., 2008) while β1 antagonism improves cognition (Ramos et al., 2005). Propranolol is a nonspecific β antagonist that has a high affinity for β1 and β2 but not β3 receptors(Schena & Caplan, 2019). Previous studies found that when administered systemically or intra-PFC, propranolol had no effect on a working memory task (Arnsten & Goldman-Rakic, 1985; B.-M. Li & Mei, 1994) suggesting that the behavioral deficits found here are not due to working memory. Together this indicates that the behavioral and neural deficits found here after propranolol are not impaired memory, but rather direct deficits in decision execution due to a disruption in representation of goal-directed action plans in M2.

β adrenoreceptors are found throughout the rat brain. In the cortex both β1 and β2 receptors are heavily expressed while there is only limited β3 expression (Nicholas et al., 1991; Rainbow et al., 1984; Schena & Caplan, 2019; Wanaka et al., 1989). Within M2 we identified a sex difference in adrenoreceptor distribution with higher densities of β2 mRNA in females (Figure 5D). Our data suggest that propranolol, acting on these β receptors, decreased suppression of irrelevant task information, reducing the signal to noise in the neural representation of task-related actions. This effect was most pronounced in females, indicating a greater reliance on adrenergic regulation of action planning in females. After propranolol, M2 neurons had limited ability to create and maintain appropriate action plans prior to decision execution, as seen by the reduction of preferred and non-preferred population activity in females (Figures 3L &4L), and reductions non-preferred activity in males (Figures 3M & 4M). Despite sex differences in β adrenergic expression, males were not immune to the behavioral effects of altered neural activity after β receptor antagonism. Both males and females had rightward shifts in decision time distributions. Overall, these behavioral deficits were mediated by β adrenergic dependent alterations in decision related activity in M2. While both males and females display neural and behavioral deficits after propranolol administration, the larger impact on female behavioral performance may be attributed to greater neural disruption after propranolol, in part due to greater dependence on β adrenoreceptors in M2.

Some aspects of decision-making strategies have known sex differences (Chen, Ebitz, et al., 2021; Chen, Knep, et al., 2021; Orsini & Setlow, 2017; Shansky, 2018) and the LC-NE system has known sex differences in its structure, size, and stress susceptibility (Bangasser et al., 2011; Pinos et al., 2001; Valentino et al., 2012). We have identified sex differences in decision-making and M2 function with regard to their sensitivity to perturbations of noradrenergic tone. We also identified a potential mechanism for this differential sensitivity by showing that within M2 there are distinct differences in β adrenergic receptor expression density. Females showed a higher density of β adrenergic receptors in M2, specifically β1 receptors on glutamatergic neurons and β2 receptors on GABAergic neurons (Figure 5). Prior evidence has demonstrated that females are more risk averse in their decision making (Orsini & Setlow, 2017). Having mechanisms for enhancing GABAergic tone in decision making regions of the brain would facilitate a risk averse behavioral strategy by slowing decision times to allow greater information accumulation and reducing impulsivity. In the current study we found that the reduction in accuracy in females after propranolol was driven by an increase in omissions, rather than incorrect responses reflecting this risk averse propensity. We also saw greatly increased decision thresholds in females when the irrelevant information in M2 was no longer suppressed. Noradrenergic regulation of local GABAergic interneurons in M2 appears critical for suppression of irrelevant action plans during decision making. With increased β expression in M2 of females, our results provide a biological basis for why disrupting this signaling would results in more pronounced physiological and behavioral outcomes in females. This highlights the potential for functionally important sex differences in susceptibility to perturbations of neuromodulation.

M2 units represent task related information and actions plans, both critical for decision-making. M2 selectively enhances specific neural populations and actively suppresses others in an effort to establish and maintain an action plan. We found that blocking β adrenergic signaling led to a reduction of task relevant encoding in M2 and decreased suppression of irrelevant motor plans. The remaining neural representation of action selection in M2 showed delayed onset, reduced discriminability, decreased magnitude, and limited duration. This disruption in M2 neural activity, delayed and impaired decision-making as seen by increased decision times/thresholds, reaction times, and omitted trials. Within these impacts we saw a sex bias where females were more disrupted by blockade of β noradrenergic signaling than males during simple decision-making. We also saw that females had greater densities of β adrenoreceptor mRNA in M2 than males, providing a potential mechanism for this bias. Results from this study indicate that propranolol, by lowering β noradrenergic signaling, can impair focused decision-making and M2 task related neural activity. Further, that sex-specific interactions in noradrenergic signaling influence cognitive function. The measures of neural activity we observed indicate an important role of β noradrenergic receptors in maintenance and creation of an action plan during decision-making, specifically in inhibiting non-preferred task related activity, increasing the signal-to-noise of task relevant neural representations.

## MATERIALS AND METHODS

### Animals

Long Evans rats (total n= 27; female n=14, male n=13; 6-8 weeks old; 200-275 grams) were purchased from Charles River (Wilmington, MA) and single housed on a reverse 12h light/dark cycle (lights ON at 9:00 PM). Separate cohorts of animals were used for in situ hybridization (female n=4, male n=4) and behavioral pharmacology/electrophysiology (female n=10, male = 9). Animals had access to ad libitum chow and water until behavioral testing began and were then restricted to 80% of ad libitum chow (female=15g, male=20g). Weight was monitored throughout experiments to ensure maintenance of at least 80% free fed weight. All procedures were approved by the Institutional Animal Care and Use Committee at the University of Massachusetts Amherst in accordance with the guidelines described in the US National Institutes of Health *Guide for the Care and Use of Laboratory Animals (National Research Council 2011)*. All animals were handled by an experimenter prior to electrode implant surgery and for 2 weeks before behavioral training began. All animals were included in behavioral analysis. Of those animals, individuals with clearly identifiable single units that completed at least 20 trials during a recording session were included in electrophysiology analysis.

### Custom static electrode array fabrication

HML VG bond coated stainless-steel wire (25μm diameter, California Fine Wire Company, Grover Beach, CA) was spun and bonded into stereotrodes with a Tetrodetwister (Open Ephys, Cambridge, MA). Each array was fabricated with 16 stereotodes and 50μm reference (1mm above electrode tips) mounted to a nanostrip connector (Omnetics, Minneapolis MN). A 32G insulated copper ground (Remington Industries, Johnsburg, IL) was connected to a stainless-steel screw in the occipital plate (McMaster Carr, Elmhurst, IL). A secondary 125μm stainless steel ground wire was placed between the dura and the skull over the parietal lobe contralateral to the implant. On the day of implant, stereotrodes were electroplated with a gold-noncyanide solution (50% diluted in saline, Sifco, Independence, OH) to achieve a target impedance of 0.1-0.2 MΩ.

### Surgery

For 2 days prior and 5 days post-surgery, animals had access to minocycline treated water (100mg/L, Ohm Laboratories, North Brunswick, NJ), weighed and replaced daily to maintain freshness and monitor water consumption. Minocycline has been shown to reduce inflammation and scarring of brain tissue after electrode implantation (Rennaker et al., 2007). To reduce respiratory secretions during surgery animals were given atropine (0.04mg, IP, Henry Schein Animal, Melville, NY) 10m before anesthesia. Animals were anesthetized with vaporized isoflurane (4% induction, ~2% maintenance) and administered the analgesic/anti-inflammatory Metacam (1 mg/kg, IP, Henry Schein Animal) and antibiotic cefazolin (0.33mg, IM, McKesson, Irving TX) prior to surgery. Animals’ eyes were covered in ophthalmic ointment (Lubifresh P.M., Livonia, MI) to prevent drying, and the incision site was shaved and sterilized with iodine and 70% isopropyl alcohol. During surgery, body temperature was maintained with deltaphase thermal pads (Braintree Scientific, Braintree, MA). Animals were placed into a stereotaxic frame (David Kopf Instruments, Tujunga, CA) and the skull was leveled. A unilateral (randomized) craniotomy was made over M2 (females right n=5, left n=3; males right n=2, left n=4). Four stainless steel 0/80 screws (McMaster Carr) were inserted into the skull to act as anchors for the implant. Electrode array was slowly lowered to the coordinates (A/P: +3mm M/L: ±1.2mm D/V: −1.5mm) at a rate of 0.2mm/min. The craniotomy was packed with sterile Gelfoam (Pfizer, Kalamazoo, MI) and covered with dental cement (OrthoJet, Wheeling, IL). Metacam and cefazolin were administered 2 days post-operatively to reduce inflammation, pain, and risk of infection. Animals were monitored and allowed to fully recover for at least 2 weeks post-surgery before beginning training.

### Behavioral task

Operant chambers (Med Associates, St Albans, VT) contained a central well with a recessed panel of LED cue lights and infrared (IR) entry beam, two laterally located levers, a speaker, and a house light. A continuous fan and sound attenuating chambers muffled outside noise. Behavioral data collection was controlled with MedPC IV (Med Associates). For all training and testing rats performed for 15% sucrose reward (95μl/trial) and animals could perform up to 250 trials in a session. All training and tests were performed during the animal’s active cycle in a dimly red lit room.

Two weeks after surgery animals were food restricted and trained on a fixed response schedule (FR1) operant task until they performed >50 responses on each lever within 1h (~4 days). They then performed a single self-initiation session where animals nose poked the IR beam to receive a sucrose reward. Next, animals were shaped to associate spatially distinct LED cues with one lever for reward. Finally, animals performed a 2AFC task where rats self-initiated trials by breaking the IR entry beam to the central well during a 40-min session (max 250 trials). Following a variable hold in the well (200-700ms), either left or right LED cue light was illuminated (50% probability). Premature withdrawal from the well before cue onset terminated a trial and was punished with a 10s cued timeout. The LED cue remained illuminated until rats withdrew from the well. Correct responses on the cued lever produced 15% sucrose reward paired with a 100ms 5kHz tone followed by a 5s intertrial-interval (ITI). Incorrect responses were punished with a 10s cued timeout. Failure to respond on a lever within 5s of withdrawal from the well counted as an omission and was punished with a 10s timeout. ITIs and timeouts were indicated by house light illumination; extinguishing the house light signaled to rats that a new trial could be initiated. Rats were trained until their individual baseline performance was stable for at least 3 days based on total trials (+/-10%) and accuracy (>70% overall). When animals showed stable performance, electrophysiology recordings were obtained. On test days, 30m before testing began rats were lightly and briefly anesthetized with isoflurane for head stage attachment and drug delivery.

### Drugs

The β adrenergic antagonist propranolol ((S)-(-)-propranolol (10mg/kg), Tocris, Bristol, UK) was diluted in 0.9% saline to achieve a final concentration of 10 mg/ml. Drugs and vehicle (0.9% saline) were administered IP 30m before testing, at a volume of 1ml/kg. Drug order was randomized, and each test day was followed by 2-3 days of baseline performance.

### Data acquisition/Electrophysiology recordings

Neural recordings were digitized at the headstage (HST/32D, Plexon; Dallas TX), passed through a passive commutator and collected using an Omniplex data acquisition system (Plexon). Signals were sampled at 40kHz, bandpass filtered 0.10Hz to 7500Hz, gain 1x, and stored offline for analysis. Recordings began five minutes before and ended five minutes after the task. MedPC timestamps were integrated through the Omniplex acquisition and animal activity was observed and recorded during the task with a video camera during the session.

### Brain collection and electrode placement

At the end of behavioral experiments, animals were anesthetized with isoflurane and electrolytic current was passed through 4-6 wires per array (10μA for 10s) for posthoc verification of electrode position. Forty-eight hours later animals were anesthetized with a ketamine xylazine solution (1.5 ml/kg, ketamine:xylazine 56:8.7 mg/ml) and transcardially perfused with ice-cold 0.9% saline followed by 4% paraformaldehyde. Immediately after, animals were perfused with 3% potassium ferrocyanide, 3% potassium ferricyanide, and 10% HCl solution for a Perl’s reaction in iron deposits at the site of electrode tips. Brains were post fixed in 4% paraformaldehyde overnight and cryoprotected in 20% sucrose azide prior to freezing and sectioning at 40μm (CM3050S; Leica, Wetzlar, Germany). Tissue slices containing M2 were stained with neutral red and electrode placement was validated. Secondary validation of lesion placement was obtained by antibody staining for glial-fibrillary associated protein (GFAP).

For GFAP staining, tissue sections were washed with PBS, blocked for 60m in PBST and 3% NDS (Jackson Immuno Research, West Grove, PA) and incubated overnight on a shaker at room temperature in fresh blocking solution with polyclonal rabbit anti-GFAP (1:1000, Dako, Santa Clara, CA). The following day tissue was washed in PBST and incubated for 2.5h at room temperature with Alexa Fluor 488 donkey anti-rabbit secondary (1:500, Jackson Immuno Research). Tissue was counterstained with in DAPI (1:1000, Millipore, Burlington, MA) in PBS for 10m. Tissue slices were washed in PBS, Tris buffer, mounted onto slides (Superfrost Plus, Fisher, Waltham, MA), covered with antifadant (Citifluor AF-1, Electron Microscopy Science, Hatfield, PA) and coverslipped. Tissue was imaged using a Zeiss Axio Imager M2 microscope (Zeiss, Oberkochen, Germany).

### In situ hybridization (RNAscope)

Animals were terminally anesthetized with a ketamine xylazine solution as above. Brains were removed, flash frozen in cooled isopentane and kept at −80C until sectioning. Frontal blocks containing M2 were embedded in OCT compound (Sakura Finetek USA, Torrance, CA), and cryosectioned at 16um onto Superfrost Plus slides (Fisher) Slide mounted sections were processed using RNAscope Muliplex Fluorescent Reagent kit (Advanced Cell Diagnostics, Newark, CA) according to the manufacturers protocol with probes targeted to β-adrenergic receptors Rn-Adrb1 (468121) or Rn-Adrb2-C2 (468131-C2), and vesicular transporters for glutamate Rn-Slc17a7(317001-C3) or GABA Rn-Slc32a1(424541). Tissue was counterstained using DAPI and imaged using a Zeiss Axio Imager M2 microscope. One M2 ROI was imaged at 20x from each hemisphere in all animals and used for manual quantification with FIJI (Schindelin et al., 2012).

### Data processing, analysis, and visualization

Offline, cross channel (>80%) artifacts were removed and a lowcut 250 Hz Bessel filter was applied. Single units (>4σ above background) were manually isolated using principal components analysis in Offline Sorter (Plexon). This resulted in a total of 347 well isolated single units for analysis: 205 units (female n=98, male n=107) from 14 animals (female n=8, male n=6) after saline, and 142 units (female n=61, male n=81) across 11 animals (female n=5, male n=6) after 10mg/kg propranolol.

Electrophysiology data was analyzed using Neuroexplorer (Plexon) and MATLAB (Natick, MA). We analyzed two task epochs of interest: cue onset, ‘Cue’ and lever press after cue presentation, “Lever press’. Data from single units were Z-scored in 100ms bins based on activity across the entire session. Z-scored data is presented as the population mean ± SEM with trials separated by side preference.

Heatmaps for each sex and drug treatment indicate the p-value of t-tests at each 50ms bin for each individual neurons’ activity on right vs. left trials with white bins not significantly different (p>0.05), and yellow (p=0.05) through purple indicating a p-value approaching 0. The onset of two consecutive significantly different bins classified the preferred direction in favor of the highest average Z-score. Side preference was determined for each unit using Z-scored data with a 50s sliding window analysis around each task epoch (−1s to 3s aligned to cue onset, −3s to 1s aligned to lever press). Latency was identified as the timepoint where 2 consecutive 50ms sliding bins showed significantly different activity on left vs right trials for each individual neuron.

To detect task relevant neurons, we identified neurons with increased activity around either cue presentation or lever press response (saline: 176 out of 205 neurons, propranolol: 116 out of 142 neurons). Units were then classified for side preference (a known feature of M2) at cue presentation and lever press response. For most neurons the side preference was consistent for cue and lever press side (saline unchanged=73.30%, changed=26.71%; propranolol unchanged=80.17%, changed=19.83%). Overall, after saline administration 85 neurons were classified as right preferring (female n=39, male n=46) and 91 were left preferring (female n=37, male n=54). After propranolol administration, 57 were classified as right preferring (female n=21, male n=36) and 59 were left preferring (female n=23, male n=36). After classification, all electrophysiology analysis only included neurons with a side preference (saline=176, female n=76, male n=100; propranolol=116, female n=44, male n=72).

To compare decoding accuracy of M2 neurons representing cue type after saline vs propranolol treatment, we used a linear support vector machine classifier. We decoded cue type (right vs left) in male and female M2 neurons after saline and propranolol using the Neural Decoding Toolbox (Meyers, 2013). We sparsely trained the classifier with a random sample of 10 presentations of each cue (right, left) from 40 random neurons from each population (saline female, saline male, propranolol female, propranolol male). We compared the accuracy of the decoding analysis within each group to shuffled neural data. This analysis was performed within each group saline female, saline male, propranolol female, propranolol male.

Behavioral and histological data was analyzed using GraphPad Prism v8 (San Diego, CA). Behavioral data is presented as mean ± SEM after confirming normality (Shapiro-Wilk) unless noted otherwise in the figure legend. Comparisons between groups were made with REML ANOVA and Sidaks post hoc tests. Distributions of firing rates and latency to cue and action plan encoding were compared using Kolmogorov– Smirnov tests. Distributions of decision times and reaction times were compared using Brown Forsythe tests. Population Z-scores on preferred and non-preferred trials were compared using t-tests. Graphs were compiled in Prism or MATLAB and figures were compiles in Adobe Illustrator CS (San Jose, CA).

## ACKNOWLEDGEMENTS

The authors would like to thank Michael Kelberman and Kara Conlan for their help training animals.

## DECLARATIONS OF INTEREST

The authors declare no competing interests.

## Notes

### Competing Interest Statement

The authors have declared no competing interest.

